# Allelic variation at a terpene synthase locus shapes wine monoterpene composition and aroma

**DOI:** 10.64898/2026.05.09.723628

**Authors:** Jerry Lin, Annegret Cantu, Larry Lerno, Mireia Domenech-Lopez, Olivia Haley, Hildegarde Heymann, Susan E. Ebeler, Dario Cantu

## Abstract

The genetic basis of wine monoterpene variation and its sensory consequences remain poorly understood. We investigated how allelic variation at a chromosome 10 linalool QTL harboring a terpene synthase cluster, including *VviTPS54*, influences wine aroma in a Riesling × Cabernet Sauvignon mapping population. Twenty-six wines from progeny selected for contrasting genotypes were subjected to HS-SPME-GC-MS profiling and descriptive sensory analysis. Genotypes carrying the high-linalool Riesling allele produced substantially elevated monoterpene levels dominated by (3*S*)-linalool. Berry and wine monoterpene profiles were significantly correlated, and berry monoterpene glycosides showed broad correlations with wine monoterpenes. High-monoterpene genotypes were more closely associated with Floral, Tropical Fruit, and Apricot/Peach attributes, though perception was modulated by matrix effects from berry pigmentation. These results demonstrate that allelic variation at a terpene synthase locus drives coordinated changes in wine monoterpene composition and linked sensory attributes, while underscoring the complexity of translating genetic variation into sensory outcomes.

## Introduction

Aroma is a central aspect of the wine experience and strongly influences consumer perception.^1^ Many components of wine aroma originate from compounds produced by the grape berry or are formed from grape-derived precursors during fermentation.^2,3^ Among these compounds, monoterpenes represent an important class of diverse odor-active molecules that contribute floral, citrus, and fruity notes to wine.^2,4^ Compounds such as (3*S*)-linalool, geraniol, and (-)-*cis*-rose oxide are particularly associated with aromatic cultivars such as Muscat, Riesling, and Gewürztraminer,^5–7^ though even in neutral cultivars, their low odor thresholds can influence sensory perception^8^. Variation in monoterpene composition contributes substantially to differences in wine aroma profiles across cultivars.^4,9–11^

Despite advances in identifying genetic loci associated with monoterpene accumulation in grape berries,^12–14^ the extent to which this variation translates into differences in wine aroma and sensory perception remains poorly understood. Fermentation and aging substantially alter volatile composition, complicating efforts to link berry chemistry to the sensory attributes of finished wine.^1,15–17^ While several studies have characterized berry volatile composition in intraspecific and interspecific wine grape populations, none have extended this analysis to winemaking and descriptive sensory evaluation.^18–21^

In earlier work,^13^ we profiled the berry metabolite composition of an F_1_ population derived from a cross between Riesling and Cabernet Sauvignon, two commercially important wine cultivars with distinct aroma profiles.^22^ Through quantitative trait locus (QTL) mapping and transcriptomic analyses, we identified a major QTL on chromosome 10 of the Riesling genome containing *VviTPS54*, a functionally characterized (3*S*)-linalool synthase whose expression closely tracked (3*S*)-linalool abundance across genotypes.^13,23^ A single nucleotide polymorphism (SNP) marker, CHR10_5431663, was identified as a reliable predictor of (3*S*)-linalool content in berry tissue.

Here, we investigate how allelic variation at the chromosome 10 QTL influences wine aroma chemistry and sensory perception. A subset of the RxCS population segregating for this locus was selected for experimental winemaking. Wine volatile compounds were quantified by HS-SPME-GC-MS, and the resulting wines were evaluated by descriptive analysis with a trained sensory panel. By integrating chemical and sensory data, this study asks whether genetic differences driving monoterpene accumulation in grape berries translate into perceptible differences in wine aroma.

## Materials and Methods

### Plant Materials and Sampling

This study utilized a subset of genotypes from a mapping population resulting from a cross between *Vitis vinifera ssp. vinifera* cv. Riesling clone FPS 24/GM 110 and *V. v. vinifera* cv. Cabernet Sauvignon clone FPS 08 (RxCS). This population was planted at the Viticulture and Enology Department Experimental Station (Oakville, Napa, CA) on *V. rupestris × riparia* Couderc 3309 rootstock in 2018 as three randomized complete blocks with 9 replicates of each genotype. A border consisting of a minimum of 6 vines of Cabernet Sauvignon clone 08 surrounded the experimental vines in order to minimize edge effects. The 140 genotypes (Riesling clone 24, Cabernet Sauvignon clone 08, and 138 progeny) in this vineyard were trained to bilateral cordons and pruned to two bud spurs. The canopy was trained in a vertical-shoot-position system modified with a crossarm with rows in a northwest-southeast orientation. Vine spacing was 1.8 meters with rows 2.4 meters apart. This population has been previously used for identifying quantitative trait loci (QTLs) associated with metabolite production, cluster architecture, and microbiome composition.^13,24,25^ A major QTL located on chromosome 10 linked to (3*S*)-linalool production was identified from this population. The marker CHR10_5431663 located in the QTL was used for the selection of genotypes for this study. In 2021, 24 RxCS genotypes were chosen for winemaking and descriptive analysis alongside the parent cultivars. Genotypes were selected using the following criteria: sufficient yield to produce enough wine for all replicates and training samples for descriptive analysis, equal numbers of pigmented and unpigmented genotypes, and equal numbers of TT and CT genotypes at marker CHR10_5431663. Berry sampling was initiated 12 weeks after flowering. A composite sample was taken from the three replicates of each experimental genotype in one block of the trial. Due to the limited amount of fruit, each sample consisted of 10-20 berries collected from different locations on multiple clusters. Samples were then crushed by hand. Juice Brix was measured weekly using a refractometer until 14 weeks post-bloom, increasing to biweekly thereafter until harvest.

### Winemaking

Grapes were hand harvested for winemaking when sugar levels reached 21 Brix or stabilized for three consecutive sampling dates. Due to low yield, fruit from all nine biological replicates in all three blocks was combined for each genotype. Following mechanical destemming and crushing, the must was adjusted if needed to a titratable acidity (TA) of 6 g/L and yeast assimilable nitrogen (YAN) of 250 mg/L using tartaric acid and diammonium phosphate, respectively. A 15% potassium metabisulfite solution was added at crush to achieve 50 mg/L of SO_2_. All fermentations were conducted in 22-liter plastic buckets at 18 °C and inoculated with 0.2 g/L of *Saccharomyces cerevisiae* strain EC1118 (Lallemand, Montreal, Canada) according to manufacturer directions. Manual punch downs were performed twice daily until fermentation was complete. Completion of fermentation was determined by a residual sugar of less than 2 g/L. The wines were then hand-pressed into 4- or 8-liter glass carboys and allowed to settle for three days. Wines were then racked off lees, sulfite adjusted to 50 mg/L SO_2_, and stored at 4 °C until bottling. Any headspace was sparged with N_2_ gas. The wines were bottled by hand after five months into 375 mL clear glass bottles and stored at 14°C until sensory and chemical analysis.

### HS-SPME-GC-MS Analysis of Wine Samples

Head-space solid-phase microextraction gas chromatography mass spectrometry (HS-SPME-GC-MS) was used to analyze volatile compounds in the wines following the protocol from Hendrickson et al.^26^ Berry volatile compound and monoterpene glycoside data for a subset of genotypes were taken from a previous study.^13^ Wine samples for volatile analysis were collected at the time of sensory evaluation and stored at 4 °C for no more than one month prior to analysis. Samples (10 mL) were placed in 20 mL amber glass headspace vials (Agilent Technologies) with 3 g NaCl and 50 µL of a 10 mg/L undecan-2-one solution (internal standard; Sigma-Aldrich, St. Louis, MO, USA), sealed with PTFE/silicone septa, and vortexed prior to analysis. Extraction was performed using a 1 cm Divinylbenzene/Carboxen/Polydimethylsiloxane (DVB/CAR/PDMS) SPME fiber (Supelco, St. Louis, MO, USA). Samples were equilibrated at 30 °C for 5 min with agitation (500 rpm), followed by fiber exposure for 45 min at 30 °C with agitation (250 rpm). The fiber was desorbed in the GC inlet at 260 °C in split mode (10:1). Chromatographic separation was carried out on an Agilent 7890A GC system coupled to a 5975C inert XL MSD and equipped with a Gerstel MPS2 autosampler. Separation was achieved using a DB-WaxETR capillary column (30 m × 0.25 mm × 0.25 µm; Agilent Technologies) under constant pressure with helium as the carrier gas and the GC method was retention time locked based on undecan-2-one.^26^ The oven temperature program was as follows: initial temperature of 40 °C held for 5 min, ramped at 3 °C/min to 180 °C, then at 30 °C/min to 260 °C and held for 7.67 min (total run time 62 min). The MS operated in scan/SIM mode over an *m/z* range of 40–300 with a solvent delay of 2 min. Peak areas were normalized to the undecan-2-one internal standard, and results are therefore reported as semiquantitative relative abundances. Due to limited fruit yield, all biological replicates were combined into a single fermentation per genotype; consequently, analytical replication reflects technical rather than biological variation. Each wine was analyzed in triplicate from a single bottle. For the parental cultivars (Riesling and Cabernet Sauvignon), bottle-to-bottle variation was assessed by analyzing three bottles in triplicate each (n = 9); variation among bottles was low (average coefficient of variance < 10%).

### Descriptive Sensory Analysis

To evaluate the sensory impact of the CHR10_5431663 alleles, 26 wines (24 progeny and the two parents) were subjected to descriptive sensory analysis (DA) of wine aroma approximately three months after bottling, following the methodology outlined by Lawless and Heymann.^27^ Analysis was conducted in the J. Lohr Wine Sensory Room at the University of California, Davis, CA, with 11 panelists (5 females and 6 males between the ages of 21 and 70) recruited from the university and Davis, CA community. Approval from the university Institutional Review Board (UCD IRB; IRB project number 1867764-1) and informed consent from the panelists were obtained prior to the onset of the study. Panelists were not cognizant of the purpose of the study. The panelists were trained over ten one-hour training sessions through which a consensus list of attributes was generated. Panelists were asked to assess the aroma of the research wines blindly and generate a list of aroma attributes. This list was narrowed through group discussion. Aroma standards were prepared for each attribute on the consensus list and used to calibrate the panelists prior to each evaluation session (**Table 1**). The wines were evaluated over a two-week period and presented as a randomized complete block design with three replicates. Assessment of the wines occurred in isolated booths with positive airflow. 20 mL of wine was served in black glasses (Riedel, Sommeliers Blind Tasting glass) were used to prevent visual bias from wine color. The research wines were evaluated by rating each attribute on an unstructured line scale anchored from ‘not present’ to ‘very intense’ using Compusense® software (Compusense Inc., Guelph, ON, Canada).

**Table 1.** Aroma reference standards used for sensory evaluation of Riesling × Cabernet Sauvignon wines.

| Aroma | Reference Standard |
| --- | --- |
| Alcohol | 20 mL vodka (Ketel One Vodka) |
| Apple | 1 wedge of sliced Granny Smith apple |
| Apple Cider | 20 mL apple cider (Martinelli's) |
| Apricot/Peach | 10 mL each apricot and peach juice (Kerns) |
| Asparagus | 2 mL brine from canned asparagus (Green Giant) |
| Banana | 1 slice of banana |
| Bubble Gum | 1/12<sup>th</sup> of a piece of Hubba Bubba gum |
| Caramel | 1 tsp caramel topping (Smucker's) |
| Cheesy | 1 tsp grated Parmigiano Reggiano cheese |
| Citrus | ¼ wedge of lemon |
| Dark Chocolate | ½ tsp 100% cocoa powder (Ghirardelli) |
| Dark Fruit | ½ tsp blackberry syrup (Monin), 1 tsp blackcurrant juice (Ribena),<br>1 tbsp cherry juice (Knudsen) |
| Earthy/Musty | 1 tbsp soil (Miracle-Gro Potting Mix) and 10 mL water |
| Floral | 1% (3S)-linalool in neutral white wine (Bota box, Chardonnay) |
| Grape Juice | 15 mL Concord grape juice (Knudsen, Organic Grape) |
| Grass | ½ tbsp grass |
| Green Bell Pepper | 2 x1 cm piece of green bell pepper |
| Green Olives | 1 green olive with 3 mL brine (Mezzetta) |
| Leather | 6 1 cm strips of leather (Kiwi Brand shoelaces) with 2 mL water |
| Molasses | 1 tsp molasses (Brer Rabbit) |
| Red Fruit | ½ tsp strawberry jam, 1 tsp raspberry jam (A natural fruit spread,<br>Mountain Fruit Company) |
| Rubber | 10 thin slices rubber (ACE rubber gasket) |
| Solvent/Acetone | 10% acetone in water |
| Tropical Fruit | 1/6 piece each of pineapple, mango, and kiwi candy (HiChew) |
| Violet | 1.5 mL water with 1.5 mL violet syrup (Monin) |
| Wax/Crayon | 2 1 cm pieces of crayon (Crayola) |

### Statistical Analysis

All statistical analyses were conducted in R (R Foundation for Statistical Computing, Vienna, Austria). Data handling and visualization were performed using the tidyverse v. 2.0 suite of packages, and multivariate and univariate statistical analyses were carried out using base R (stats v. 3.6.2) and supporting packages as listed below. The potential influence of wine matrix parameters was assessed to determine whether they should be included as covariates in subsequent analyses. The effects of wine alcohol content and pH on volatile compound abundance and sensory attributes were evaluated using Spearman rank correlations (cor.test, method = “spearman”), linear models (lm), and multivariate analysis of variance (MANOVA) applied to genotype-averaged data. For MANOVA, all sensory attributes were included as a multivariate response, with alcohol content and pH treated as continuous predictors and significance assessed using Wilks’ λ. P-values from univariate tests were adjusted for multiple testing using the false discovery rate (FDR) method (p.adjust, method = “fdr”). As no significant associations were detected (P-adj > 0.05), alcohol and pH were not included as covariates in subsequent analyses. For chemical and sensory datasets, multivariate analysis of variance (MANOVA) was then used to assess overall product effects. Models for descriptive analysis (DA) data included product, judge, and replicate as fixed effects to account for panelist variability and repeated measurements. Wilks’ λ was used as the test statistic, with approximate F-tests used to determine significance. For volatile compound datasets, MANOVA models included product and replicate effects. Following MANOVA, univariate analyses of variance (ANOVA) were performed for individual variables using the same model structure. For DA data, ANOVA models included product, judge, and replicate effects, whereas chemical data models included product and replicate. P-values were adjusted for multiple testing using the false discovery rate (FDR) method (p.adjust, method = “fdr”). Attributes or compounds showing significant product effects (α = 0.05) were retained for subsequent multivariate analyses. Permutational multivariate analysis of variance (PERMANOVA) was used to evaluate multivariate differences in wine volatile profiles and assess the potential influence of wine matrix variables. Analyses were conducted using the adonis2 function in the vegan package (v. 2.7-2) ^28^. To test whether volatile composition differed among genetic marker classes, a Euclidean distance matrix was constructed from the scaled compound data and modeled as a function of marker class using 999 permutations. Separately, the effects of wine alcohol and pH on volatile composition were evaluated with alcohol and pH as explanatory variables. Principal component analysis (PCA) was performed on mean-centered, scaled chemical data using the *FactoMineR* v. 2.13 package ^29^ to visualize patterns of variation among wines. Canonical variate analysis (CVA) was conducted using the package candisc v 1.10 ^30^ on significant sensory attributes to maximize separation among products relative to within-group variation. For CVA, replicate observations were averaged within each judge × product combination to reduce within-panel noise while preserving judge-level variation. Wine centroids were calculated as the mean position across judges, and score and loading plots were used to interpret multivariate structure. Relationships between chemical composition and sensory attributes were explored using multiple factor analysis (MFA) with the package *FactoMineR* v. 2.13 to identify associations between volatile compounds and descriptive sensory attributes and to visualize their joint contribution to sample differentiation. Significance for all tests was determined at α = 0.05 unless otherwise stated.

## Results

### Genotype selection and winemaking

Metabolite profiling of berries revealed significant variation in monoterpene content between genotypes with differing alleles at the CHR10_5431663 marker (**Fig. 1**). As both parent cultivars are grown for commercial wine production, we selected a subset of genotypes for winemaking and descriptive analysis to assess the sensory impact of allelic variation at this locus. Genotype selection was constrained by the need for adequate fruit yield. Alongside both parents, 24 progeny were selected: 12 genotypes heterozygous (CT) at the CHR10_5431663 locus (6 red and 6 white), which produced moderate to high levels of (3*S*)-linalool, and 12 homozygous (TT) (7 red and 5 white), with low (3*S*)-linalool production.

**Figure 1.**
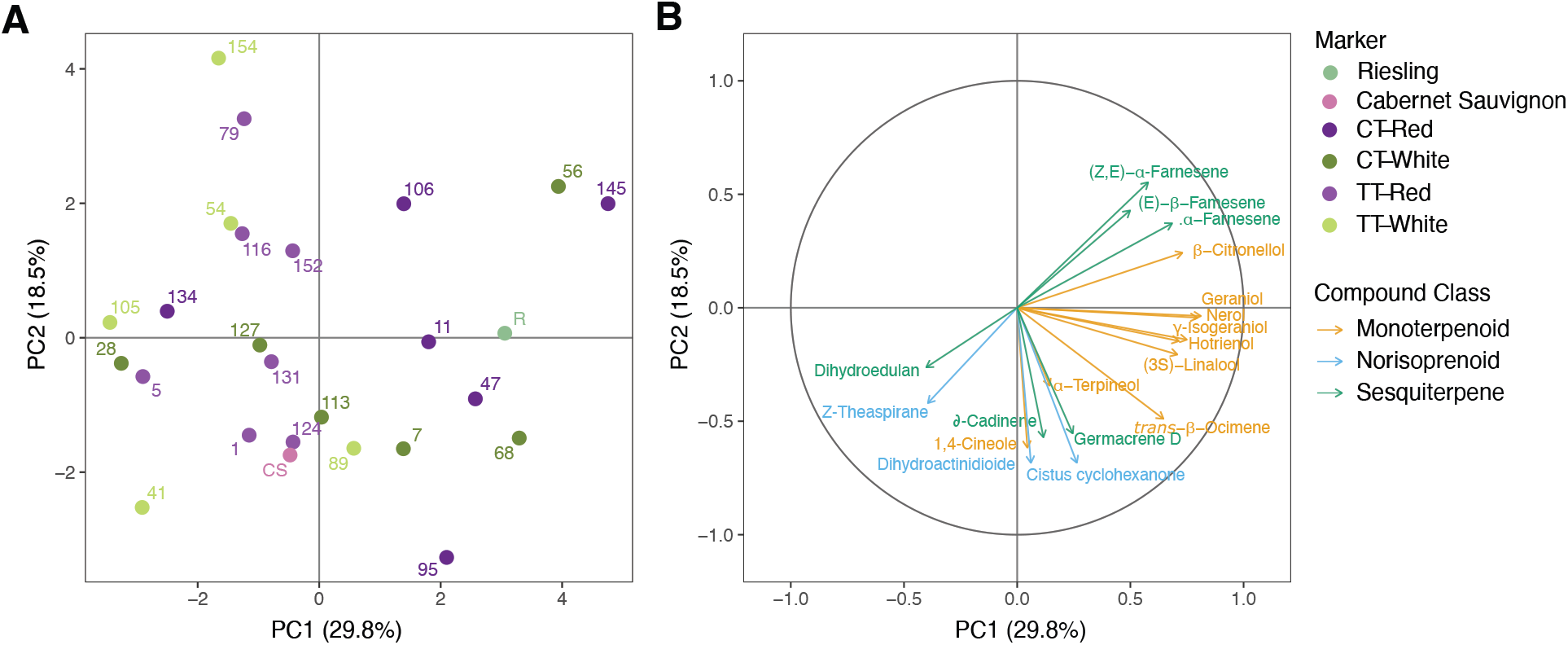
Principal component analysis (PCA) of selected RxCS genotypes used for winemaking based on berry terpene profiles, including monoterpenes, sesquiterpenes, and norisoprenoids. (A) PCA scores plot showing the distribution of genotypes colored by marker genotype class, with parental cultivars (Cabernet Sauvignon and Riesling) shown separately. (B) Corresponding loading plot showing compound contributions to PC1 and PC2, colored by chemical class.

Due to variation in phenological stages, harvest of the RxCS genotypes spanned two weeks (**Table 2**). In nine genotypes, sugar accumulation slowed or stopped before the target of 21 ºBrix was reached, and these were harvested early to avoid dehydration, sour rot, and bird damage. Consequently, ºBrix at crushing ranged from 19.1 to 24.0, with white genotypes averaging slightly higher than red genotypes (22.1 vs 21.1 ºBrix). Fruit yield ranged from 4.28 kg to 13.77 kg with a mean of 7.45 kg. All fermentations proceeded to dryness without detectable issues. Despite substantial variation in alcohol content and pH among finished wines, neither parameter significantly influenced volatile composition or sensory attributes (MANOVA, P-adj > 0.05), indicating that differences in wine chemistry and aroma were not driven by these matrix variables and were therefore excluded from subsequent analyses.

**Table 2.** Harvest parameters and resulting wine composition for selected genotypes of the RxCS population, including parental cultivars (Cabernet Sauvignon [CS] and Riesling [R]). Measured variables include fruit metrics at harvest (fruit weight, °Brix, pH, titratable acidity [TA], malic acid, and yeast assimilable nitrogen [YAN]) and finished wine properties (residual sugar, final pH, TA, alcohol content, and final volume). Genotypes are categorized by marker genotype (CT, TT) and berry color (red, white).

| Genotype | Marker Genotype | Color | Harvest Date | Fruit Weight (kg) | Brix | Initial pH | Initial TA (g/L) | Initial Malic (mg/L) | Initial YAN (mg/L) | Residual Sugar (g/L) | Final pH | Final TA (g/L) | Alcohol % | Final Volume (L) |
| --- | --- | --- | --- | --- | --- | --- | --- | --- | --- | --- | --- | --- | --- | --- |
| 1 | TT | Red | 8-Sep | 6.69 | 21.70 | 3.12 | 7.05 | 1741 | 38 | 0.00 | 3.49 | 6.9 | 12.4 | 3.79 |
| 5 | TT | Red | 27-Aug | 5.36 | 21.1 | 3.06 | 8.37 | 1862 | 122 | 0.01 | 3.31 | 7.5 | 11.6 | 3.79 |
| 7 | CT | White | 2-Sep | 5.71 | 21.6 | 3.16 | 6.36 | 1214 | 50 | 0.02 | 3.16 | 8.1 | 11.2 | 3.03 |
| 11 | CT | Red | 8-Sep | 5.81 | 20.50 | 3.23 | 5.02 | 1001 | 35 | 0.01 | 3.30 | 7.5 | 11.7 | 3.41 |
| 28 | CT | White | 2-Sep | 6.00 | 23.9 | 2.75 | 10.42 | 1913 | 65 | 0.00 | 3.44 | 7.0 | 12.1 | 3.41 |
| 41 | TT | White | 27-Aug | 6.37 | 22.4 | 3.03 | 9.06 | 3690 | 106 | 0.00 | 3.54 | 6.9 | 12.1 | 3.79 |
| 47 | CT | Red | 2-Sep | 8.23 | 20.6 | 3.45 | 5.25 | 2320 | 45 | 0.00 | 3.53 | 6.7 | 11.7 | 4.43 |
| 54 | TT | White | 9-Sep | 9.63 | 22.90 | 3.19 | 5.53 | 1418 | 98 | 0.00 | 3.26 | 7.9 | 12.3 | 5.68 |
| 56 | CT | White | 8-Sep | 8.44 | 20.60 | 3.30 | 5.15 | 1608 | 92 | 0.01 | 3.08 | 9.4 | 13.1 | 5.07 |
| 68 | CT | White | 27-Aug | 9.16 | 23 | 3.09 | 8.76 | 3904 | 168 | 0.02 | 3.24 | 8.5 | 12.8 | 5.37 |
| 79 | TT | Red | 8-Sep | 7.20 | 20.10 | 3.02 | 7.95 | 1648 | 146 | 0.02 | 3.41 | 7.5 | 12.8 | 5.07 |
| 89 | TT | White | 2-Sep | 5.85 | 21.4 | 3.00 | 7.27 | 1775 | 58 | 0.01 | 3.49 | 6.9 | 12.5 | 3.41 |
| 95 | CT | Red | 9-Sep | 6.30 | 20.60 | 3.38 | 4.87 | 1171 | 87 | 0.00 | 3.52 | 6.5 | 12.1 | 3.79 |
| 105 | TT | White | 27-Aug | 8.07 | 22.3 | 3.17 | 6.73 | 2130 | 101 | 0.01 | 3.39 | 6.3 | 12.5 | 5.07 |
| 106 | CT | Red | 2-Sep | 11.04 | 20.6 | 3.11 | 6.43 | 1498 | 64 | 0.02 | 3.28 | 6.2 | 12.8 | 5.56 |
| 113 | CT | White | 27-Aug | 7.71 | 23 | 3.19 | 6.19 | 1360 | 211 | 0.02 | 3.27 | 6.3 | 12.4 | 5.07 |
| 116 | TT | Red | 8-Sep | 10.15 | 22.20 | 3.32 | 5.68 | 2159 | 128 | 0.02 | 3.38 | 6.0 | 12.3 | 6.62 |
| 124 | TT | Red | 9-Sep | 7.58 | 21.30 | 3.29 | 5.31 | 1302 | 98 | 0.01 | 3.41 | 6.8 | 12.7 | 5.68 |
| 127 | CT | White | 2-Sep | 4.80 | 21.4 | 3.08 | 6.98 | 1362 | 51 | 0.00 | 3.33 | 8.1 | 13.2 | 3.03 |
| 131 | TT | Red | 9-Sep | 8.72 | 19.10 | 3.30 | 5.32 | 1306 | 67 | 0.00 | 3.28 | 7.6 | 12.4 | 5.68 |
| 134 | CT | Red | 9-Sep | 5.74 | 20.50 | 3.33 | 5.57 | 1756 | 97 | 0.00 | 3.23 | 7.0 | 11.3 | 4.16 |
| 145 | CT | Red | 8-Sep | 4.28 | 21.50 | 3.36 | 5.52 | 1963 | 77 | 0.01 | 3.27 | 7.1 | 11.5 | 3.79 |
| 152 | TT | Red | 2-Sep | 5.77 | 21.7 | 3.34 | 5.52 | 1956 | 139 | 0.01 | 3.30 | 7.2 | 12.0 | 3.41 |
| 154 | TT | White | 8-Sep | 6.84 | 21.60 | 3.39 | 4.94 | 1956 | 130 | 0.01 | 3.49 | 6.4 | 11.9 | 5.07 |
| CS | TT | Red | 9-Sep | 13.77 | 24.00 | 3.27 | 5.47 | 2257 | 116 | 0.01 | 3.59 | 5.8 | 11.6 | 7.57 |
| R | CT | White | 27-Aug | 8.48 | 20.7 | 3.05 | 7.1 | 1719 | 69 | 0.01 | 3.46 | 6.4 | 12.1 | 5.68 |

### Wine volatile composition varies with marker genotype

Genotypes showed substantial differences in volatile composition after fermentation (**Fig. 2** and **Fig. 3**). A total of 47 compounds were identified from the wines (**Supplementary Table S1**). PCA of all volatile compounds (**Fig. 2A)** showed that the first two dimensions explained 23.3% and 18.6% of the variance, respectively. Separation between Cabernet Sauvignon and Riesling wines was driven mainly by esters such as ethyl butanoate and phenylethyl acetate, the norisoprenoids 1,1,6-trimethyl-1,2-dihydronaphthalene (TDN) and β-cyclocitral, and monoterpenoids including (3*S*)-linalool. Progeny wines were distributed between the parents, with nonpigmented CT genotypes largely occupying the upper right quadrant.

**Figure 2.**
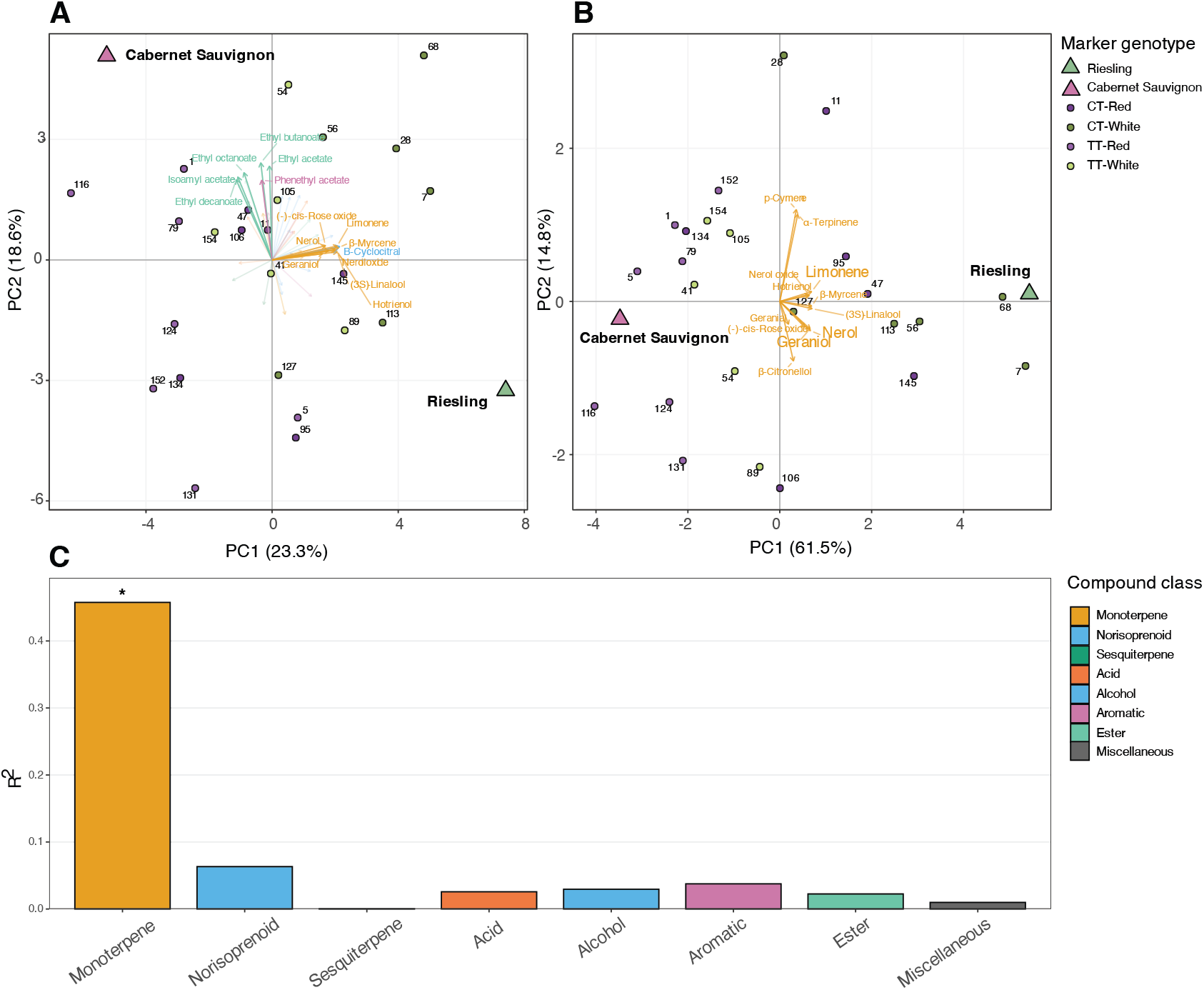
Multivariate analysis of wine volatile composition in the RxCS population. (A) PCA biplot of all detected volatile compounds. (B) PCA biplot of monoterpenes only. In both panels, points represent individual genotypes colored by marker genotype class, with parental cultivars shown separately. Vectors represent compound loadings colored by chemical class. (C) Proportion of variance in each volatile compound class explained by marker genotype (R^2^), as determined by PERMANOVA. The bar marked with an asterisk (*) indicates significance at P-adj < 0.05.

**Figure 3.**
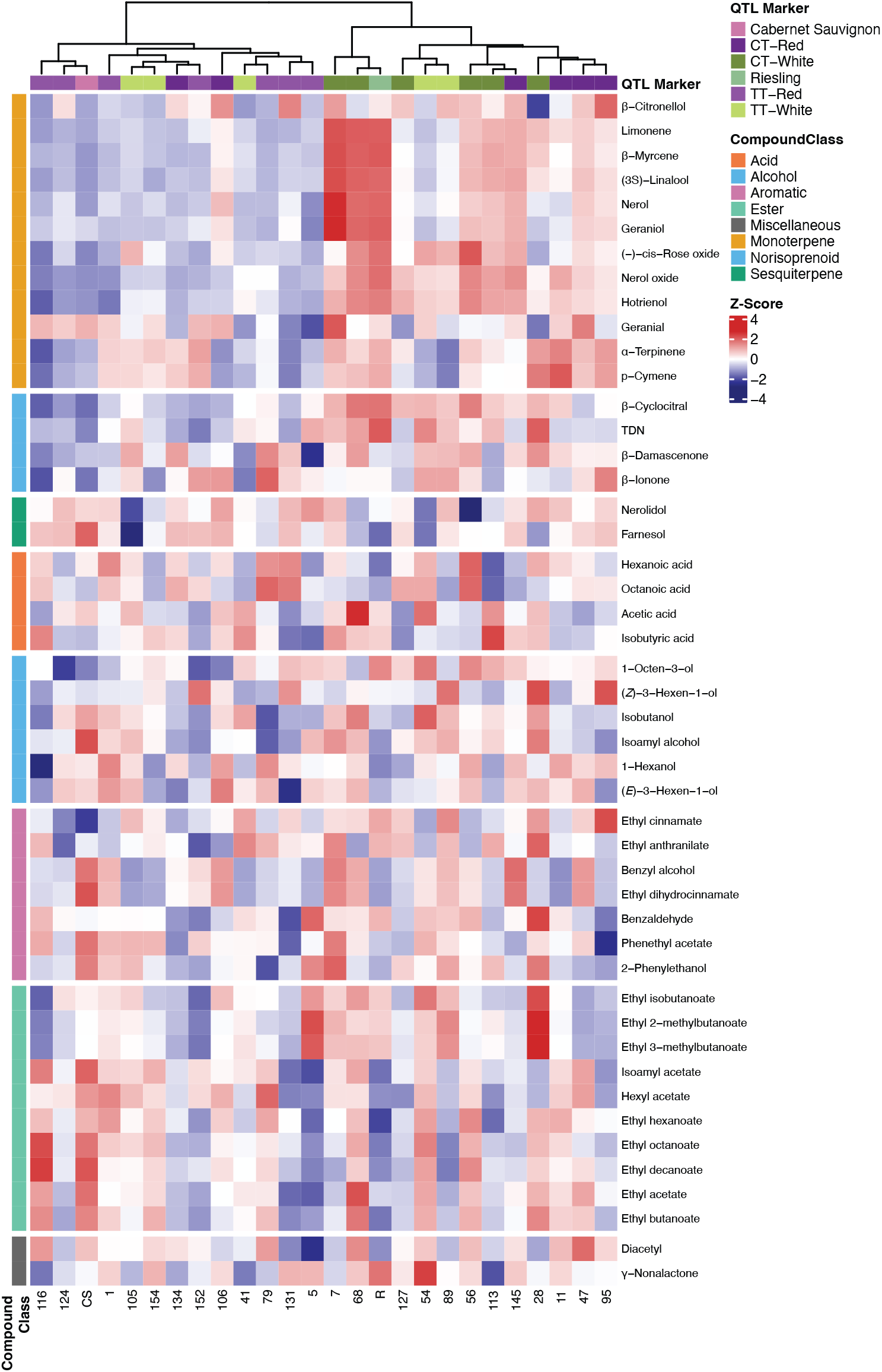
Heatmap of wine volatile compound relative abundances across the RxCS population. Compounds (rows) were hierarchically clustered using Euclidean distance and complete linkage within compound classes. Genotypes (columns) were clustered in the same manner according to monoterpene relative abundances. Values were log2-transformed and rescaled as Z-scores prior to plotting (red = higher; blue = lower relative abundance). Row annotations indicate compound chemical class. Column annotations indicate berry color (pigmented/unpigmented) and CHR10_5431663 marker genotype (CT/TT). Compound classes are separated by horizontal breaks.

Restricting the analysis to monoterpenes (**Fig. 2B**) increased the variance explained by PC1 to 61.5% and sharpened the separation between marker genotypes. Genotypes were distributed along PC1 according to (3*S*)-linalool content and that of correlated monoterpenoids. PC2 captured stratification associated with the relative abundance of β-citronellol versus the monocyclic terpenes *p*-cymene and α-terpinene, which were predominant in genotypes 28 and 11, whereas β-citronellol was more abundant in genotypes 95 and 106. PERMANOVA confirmed that the effect of marker genotype on volatile composition was specific to monoterpenes, which showed strong differentiation between TT and CT genotypes (R^2^ = 0.46, P-adj = 0.008; **Fig. 2C**).

Hierarchical clustering (**Fig. 3)** reflected this differentiation, though imperfectly: two TT individuals clustered with CT genotypes and vice-versa. While marker effects were significant only for monoterpenes (**Fig. 2C**), norisoprenoids showed notable variation among genotypes, with β-cyclocitral, TDN, β-ionone, and β-damascenone occurring at substantially higher levels in some specific individuals. TDN was most abundant in wine from Riesling, genotype 28, and genotype 54, but was detectable in all wines including Cabernet Sauvignon. TDN and β-cyclocitral were significantly positively correlated with each other and with most monoterpenes.

### Berry and wine monoterpene profiles are correlated

Berry volatile compounds and monoterpene glycosides of a subset of 10 genotypes analyzed in a previous study were compared with volatile compounds measured in the corresponding wines.^13^ Correlations between berry and wine compounds are shown in **Fig. 4**. Wine monoterpenes, including (3*S*)-linalool, β-myrcene, geraniol, limonene, and nerol were significantly correlated with a broad range of berry monoterpenes and monoterpene glycosides (r > 0.7, P-adj <0.05), indicating that berry monoterpene composition is a reliable predictor of wine monoterpene content. Correlations between the same compound in berry and wine were significant for (3*S*)-linalool and nerol, suggesting that these compounds are transferred relatively directly from berry to wine with limited transformation. Most monoterpene glycosides showed broad correlations with many wine monoterpenes, with the exception of PH1 and PH2, which were only significantly correlated with α-terpinene (P-adj <0.05) and showed a trend toward correlation with *p*-cymene (P-adj <0.1), consistent with their potential role as precursors to these monocyclic terpenes during fermentation.

**Figure 4.**
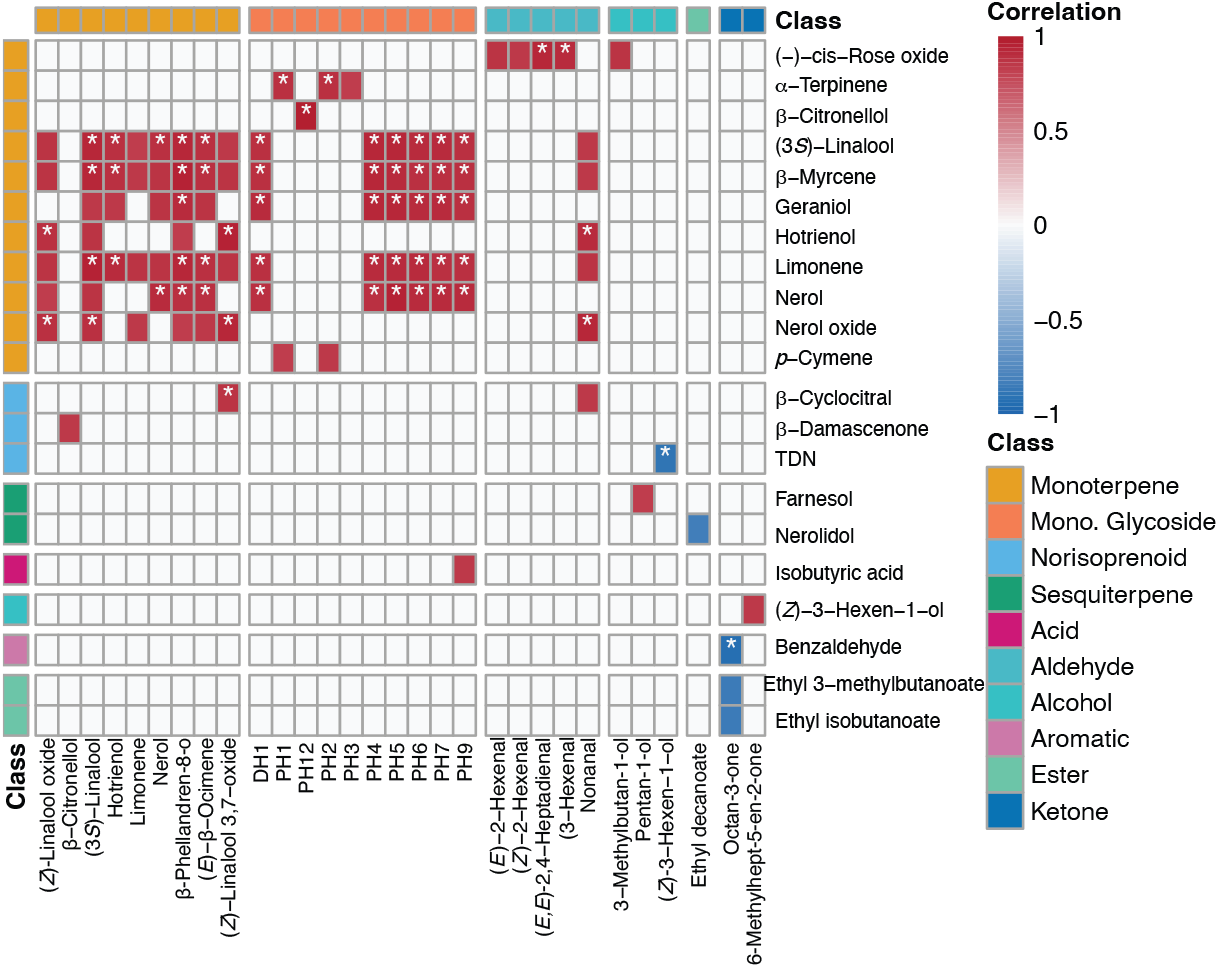
Correlation heatmap between berry-derived volatile compounds and monoterpene glycosides (columns) and wine volatile compounds (rows) in the RxCS population. Only pairs with correlation coefficients r > 0.7 and adjusted P-value < 0.1 are displayed; all other cells are blank. Color indicates the direction and strength of correlation (red = positive; blue = negative). Asterisks denote pairs significant at P-adj < 0.05. Compounds are grouped by chemical class, indicated by color-coded row and column annotations, with breaks separating major classes.

### Monoterpene composition varies continuously across the population

Wine monoterpene composition varied widely across the RxCS population, with total relative abundances spanning more than an order of magnitude among genotypes (**Fig. 5**). Monoterpene levels formed a continuous gradient, with some individuals reaching levels approaching or exceeding those of Riesling. With one exception, CT genotypes produced higher total monoterpene levels than TT genotypes; genotype 134 (CT) produced lower levels than four TT genotypes. (3*S*)-Linalool was the predominant monoterpene across all genotypes (**Table 3**), contributing 22.6-79.7% of the total, with the highest-abundance genotypes also exhibiting the greatest proportion of (3*S*)-linalool. The second most abundant wine monoterpene varied among genotypes, with β-citronellol, *p*-cymene, and hotrienol most frequently observed. Geranial was the second most abundant monoterpene in only a single genotype.

**Table 3.** Monoterpene composition of wines from selected RxCS genotypes. Total monoterpene relative abundance is expressed in undecan-2-one equivalents. The percentage contributions of (3*S*)-linalool and the second most abundant monoterpene to the total monoterpene pool are shown for each genotype. Genotypes are ordered by increasing total monoterpene abundance.

| Genotype | Marker Genotype | Berry Color | Total Monoterpenes (Relative Abundance) | (3*S*)-Linalool % Total | 2nd Major Monoterpene | % Total |
| --- | --- | --- | --- | --- | --- | --- |
| CS | TT | Red | 0.038 | 32.3 | β-Citronellol | 17.8 |
| 116 | TT | Red | 0.033 | 40.7 | β-Citronellol | 19.6 |
| 124 | TT | Red | 0.053 | 40.4 | β-Citronellol | 21.3 |
| 5 | TT | Red | 0.056 | 49.5 | β-Citronellol | 13.6 |
| 79 | TT | Red | 0.06 | 32.3 | *p*-Cymene | 16.4 |
| 131 | TT | Red | 0.066 | 43.5 | β-Citronellol | 26.1 |
| 154 | TT | White | 0.069 | 23 | *p*-Cymene | 21.1 |
| 1 | TT | Red | 0.07 | 31.8 | *p*-Cymene | 24.4 |
| 41 | TT | White | 0.075 | 42.8 | Hotrienol | 14.1 |
| 134 | CT | Red | 0.078 | 31.8 | *p*-Cymene | 24.4 |
| 105 | TT | White | 0.085 | 35.1 | *p*-Cymene | 19.7 |
| 54 | TT | White | 0.092 | 26.4 | Hotrienol | 28.9 |
| 152 | TT | Red | 0.098 | 22.6 | *p*-Cymene | 26.4 |
| 89 | TT | White | 0.117 | 41.1 | Hotrienol | 19.4 |
| 106 | CT | Red | 0.147 | 63.5 | β-Citronellol | 11.1 |
| 127 | CT | White | 0.193 | 34.5 | Hotrienol | 31.4 |
| 28 | CT | White | 0.22 | 44.4 | *p*-Cymene | 20.1 |
| 95 | CT | Red | 0.228 | 45 | *p*-Cymene | 13.5 |
| 11 | CT | Red | 0.262 | 26.8 | *p*-Cymene | 24.3 |
| 47 | CT | Red | 0.283 | 58.7 | Hotrienol | 7.2 |
| 113 | CT | White | 0.458 | 55.9 | Hotrienol | 18.5 |
| 145 | CT | Red | 0.506 | 57.1 | Hotrienol | 15.2 |
| 56 | CT | White | 0.525 | 38.8 | Hotrienol | 32.8 |
| 68 | CT | White | 1.322 | 67.4 | Hotrienol | 13 |
| 7 | CT | White | 1.696 | 79.7 | Geranial | 6.4 |
| R | CT | White | 1.357 | 60 | Hotrienol | 16.2 |

**Figure 5.**
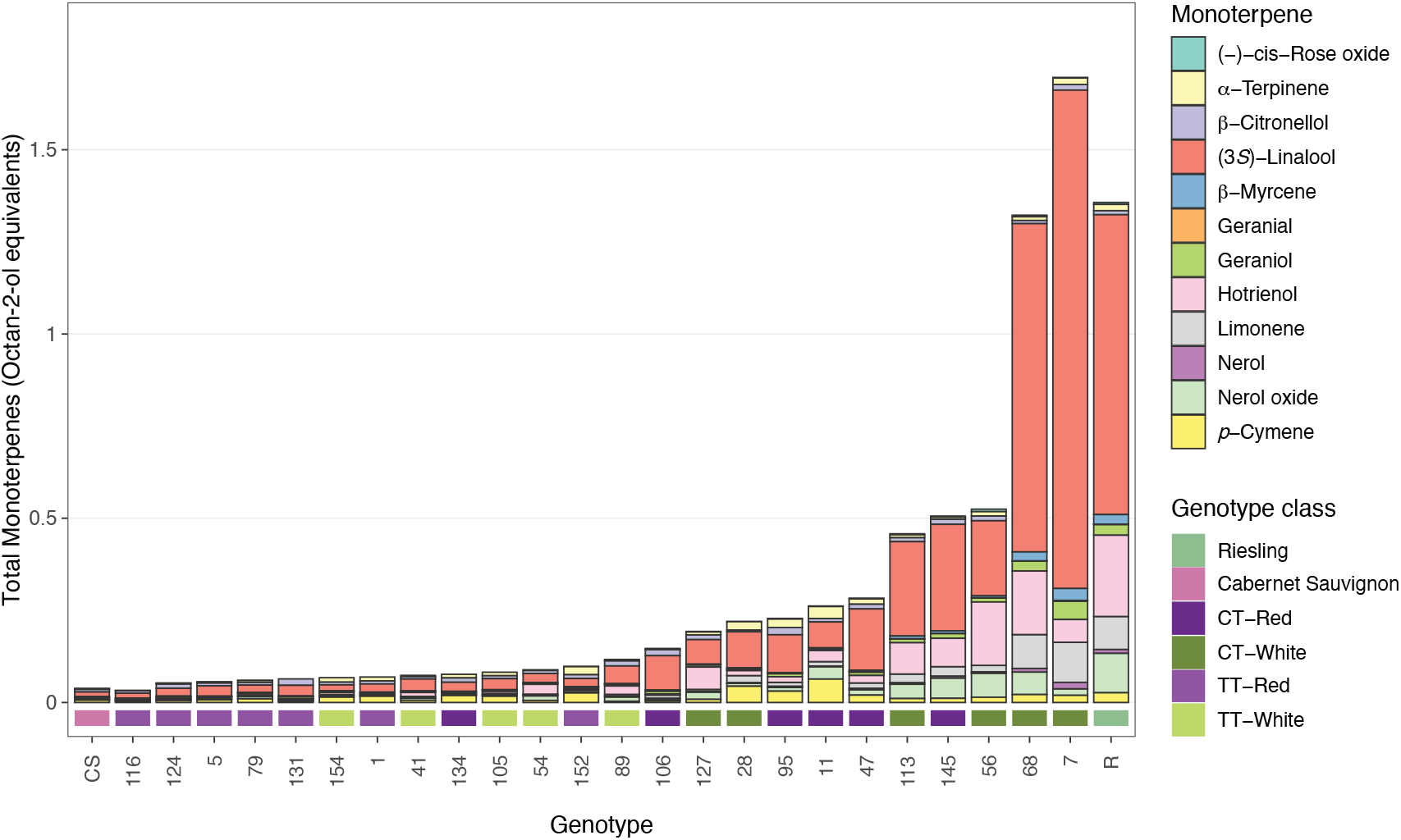
Genotype-dependent variation in wine monoterpene composition across the RxCS population. Stacked bars show the relative abundance of individual monoterpenes in wines from each genotype, expressed in undecan-2-one equivalents. Genotypes are ordered from lowest to highest total monoterpene abundance. Parental cultivars (CS, Cabernet Sauvignon; R, Riesling) are labeled separately. Genotype class is indicated by the color-coded bar below each genotype label.

### Descriptive analysis reveals genotype-dependent aroma profiles

The DA panelists generated 26 aroma descriptors by consensus (**Table 1**). MANOVA indicated significant product effects on aroma profiles in the full dataset after accounting for judge and replicate effects (Wilks’ λ = 0.13, p < 0.05, **Supplementary Table S2**). Subsequent ANOVA identified 14 attributes that were statistically significant at a P-adj of <0.05. CVA of these attributes is shown in **Fig. 6A** and **6B**. The first canonical variate (Can1) accounted for 35.2% of the variation among products, and Can2 for an additional 16.2%. Red and white genotypes were clearly separated along Can1. White genotypes showed a close association with five descriptors: Asparagus, Green Olives, Floral, Tropical Fruit, and Apricot/Peach, with genotypes 7, 28, 56, 68, 113, and Riesling showing a particularly strong positive correlation with Floral and Asparagus. Red genotypes were distinguished by three loose groupings: Dark Chocolate, Dark Fruit, Acetone, and Banana; Caramel and Cheesy; and Green, Grape Juice, and Apple. Can2 mainly separated red genotypes between the Caramel and Cheesy grouping and the other two. The aroma profiles of Cabernet Sauvignon and genotype 106 were highly driven by the Cheesy attribute. White genotypes showed less variation along Can2 than red genotypes. Within each color grouping, CT genotypes were generally located to the left of TT genotypes in canonical space.

**Figure 6.**
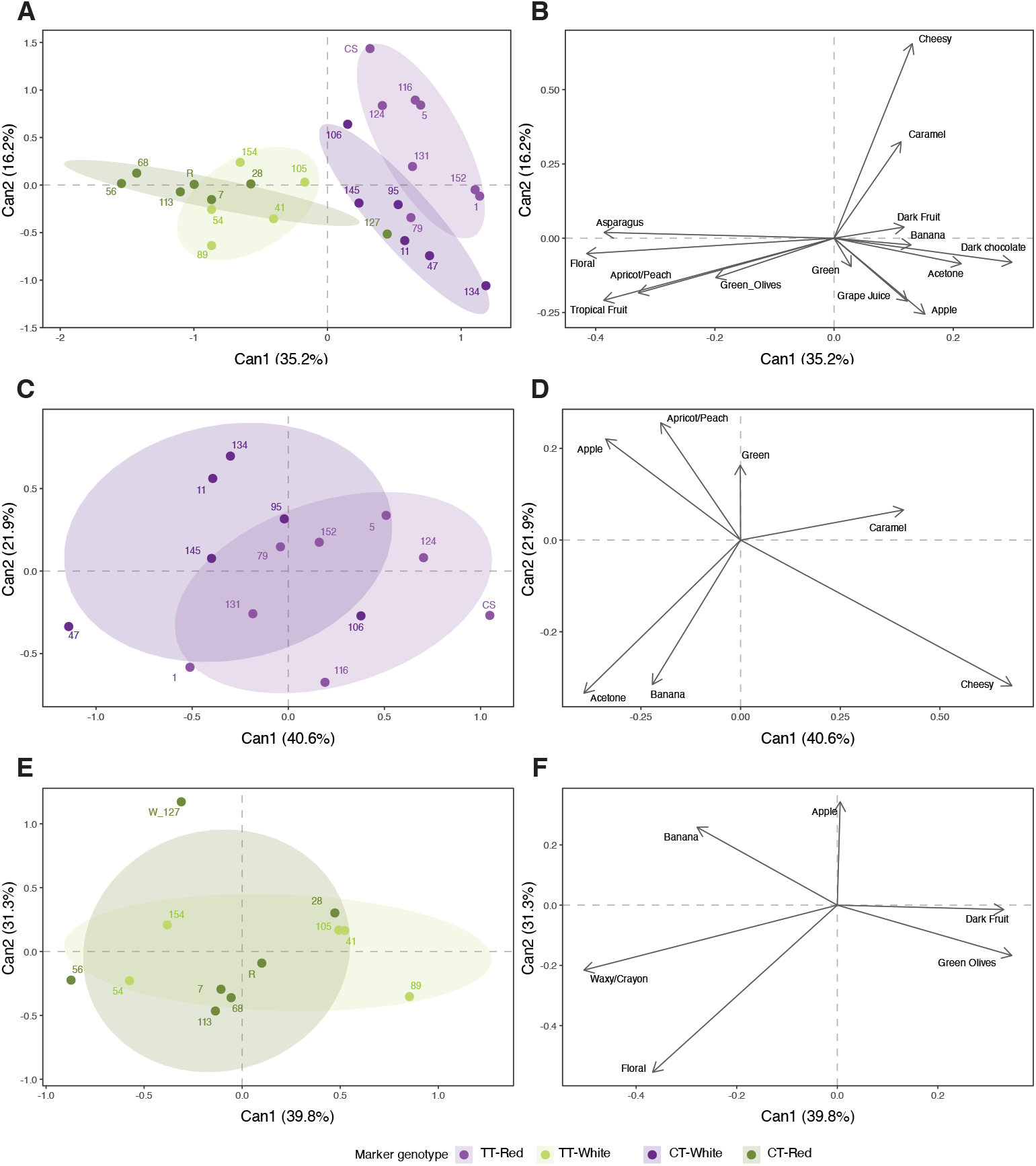
Canonical variate analysis (CVA) of descriptive analysis data from RxCS wines. (**A, C, E**) Score plots of the first two canonical variates for all wines, red wines only, and white wines only, respectively. Points represent individual wine genotypes colored by marker genotype class. Shaded ellipses indicate the dispersion of wines within each marker genotype class. (**B, D, F**) Corresponding loading plots displaying canonical structure coefficients for sensory attributes significant under ANOVA (P-adj < 0.05) within each dataset. Percent variance explained by each canonical variate is indicated on the axes. Eigenvalues are provided in **Supplementary Table S3**.

Because berry color dominated separation in the full dataset, MANOVA, ANOVA, and CVA were repeated separately on red and white wines (**Fig. 6C-F**). Product effects were significant for both subsets (Wilks’ λ = 0.20 and 0.18 for reds and whites, respectively; p < 0.05, **Supplementary Table S2**). For red wines, Can1 accounted for 40.6% of the variation (**Fig. 6C** and **D**), with wines distributed along a gradient from fruity attributes such as Banana and Apricot/Peach to Cheesy and Caramel. CT genotypes tended to score higher in Apple, Apricot/Peach, and Green than TT genotypes, though substantial overlap was present, indicating only moderate differentiation. One CT individual, 106, was an exception, showing a negative association with Apple and Apricot/Peach.

Analysis of white wines showed weaker structure (**Fig. 6E** and **F**). Can1 and Can2 explained 39.8% and 31.3% of the variation, respectively, but marker genotype classes showed substantial overlap. Nevertheless, the Floral descriptor was much more closely associated with CT individuals. The Waxy/Crayon descriptor, not significant in the full or red wine dataset, was a strong contributor to Can1 separation and positively associated with genotypes 54 and 56. Overall, marker genotype effects on aroma perception were more discernible among red wines than white wines, where genotype classes were less distinctly separated in canonical space.

### Integration of chemical and sensory data

To assess the relationship between sensory and volatile profiles, multiple factor analysis (MFA) was performed, integrating DA attributes with wine volatile compound data (**Fig. 7A-F**). As with the CVA, MFA was conducted on all wines and on subsets by berry color. When all wines were considered, the first two dimensions explained 34.8% of the variance. Dimension 1 (22.3%) distinguished genotypes primarily by berry color and secondarily by marker genotype, with white genotypes clustering on the positive side of the plot, consistent with the CVA. Within each color grouping, CT individuals were located to the right of TT genotypes, corresponding to positive loadings of monoterpenes including (3*S*)-linalool, limonene, and β-myrcene, and the sensory attributes Floral, Tropical Fruit, and Apricot/Peach (**Fig. 7B**). Interestingly, genotype 145, despite having the fourth highest monoterpene level, scored lower in monoterpene-associated descriptors than several white genotypes with substantially lower monoterpene content.

**Figure 7.**
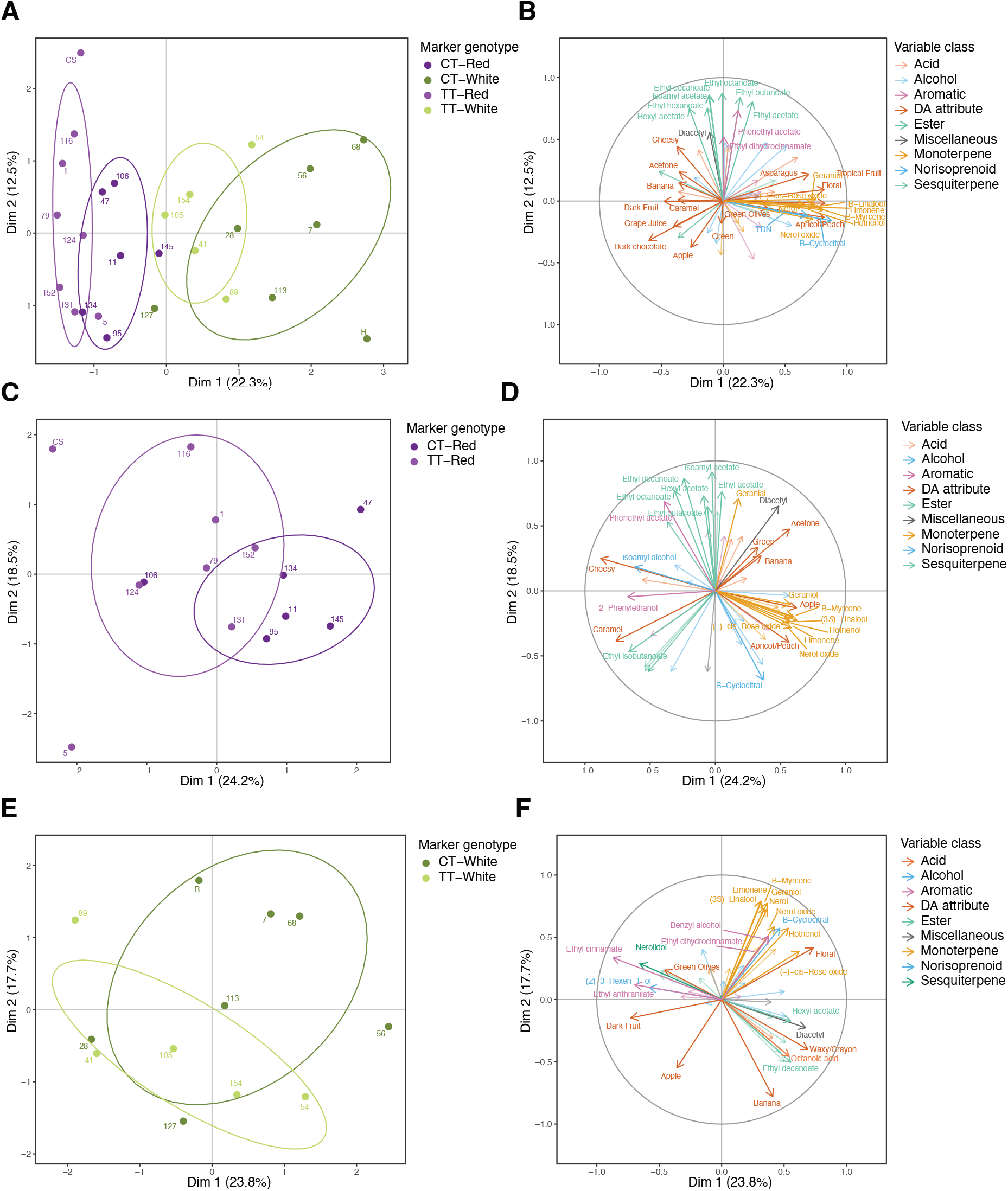
Multiple factor analysis (MFA) of wine descriptive analysis (DA) attributes and SPME-GC–MS volatile compounds across the Riesling × Cabernet Sauvignon population. **Panels A, C, and E:** MFA scores plots showing genotype clustering based on combined sensory and chemical profiles for wines produced from (A) all genotypes (n=26), (C) red genotypes (n=14), and (E) white genotypes (n=12). Points represent genotype means, colored by marker genotype class, with ellipses indicating group dispersion within one standard deviation (68% confidence interval). **Panels B, D, and F:** Corresponding variable correlation circles displaying loadings of significant DA attributes and GC–MS compounds. Arrows indicate the direction and strength of variable contributions to the first two MFA dimensions, with colors denoting compound class. DA attributes are highlighted in orange. For clarity, only the top 10 contributors to Dimensions 1 and 2 among GC–MS variables are labeled, along with all significant DA attributes. Percent variance explained by each dimension is shown on the axes. Eigenvalues are provided in **Supplementary Table S4**.

Genotypes on the left side of Dimension 1 were associated with fermentation-derived descriptors such as Cheesy, Dark Fruit, and Banana. Except for a moderate positive correlation between hexyl acetate and Cheesy, no strong associations between sensory attributes and volatile compounds were identified in this region. Dimension 2 (12.5%) was driven by the relative abundance of yeast-derived esters including isoamyl acetate, phenylethyl acetate, and ethyl hexanoate, none of which were significantly associated with any DA attributes. Of note, the fruit descriptors Banana, Dark Fruit, and Grape Juice showed little to no correlation with these esters, despite their association with fruity aromas ^17,31^.

Restricting the analysis to red genotypes (**Fig. 7C** and **D**) preserved the separation of markers along Dimension 1 (24.2%), though with reduced dispersion. Compound-descriptor associations not apparent in the full dataset emerged here: Cheesy was highly correlated with isoamyl alcohol, and Caramel with ethyl isobutanoate. Among white genotypes (**Fig. 7E** and **F**), CT genotypes largely occupied the upper-right quadrant of the plot in association with monoterpenes and the Floral descriptor, though separation from TT genotypes was incomplete. Two additional associations emerged in this subset: Waxy/Crayon, not significant in the other datasets, was correlated with ethyl decanoate and octanoic acid, both known for fatty or waxy odor character ^31,32^; and Green Olives was highly correlated with nerolidol, a sesquiterpene, and ethyl cinnamate, a phenylpropanoid-derived ester.

The association between Floral and Apricot/Peach and most monoterpenes was consistent across all three datasets (**Fig. 7B, D, F**), with increased loading separation within color subsets suggesting additional compound contributions once major between-group differences are removed. Collectively, these results indicate that genotype-dependent variation in wine aroma is driven primarily by coordinated changes in monoterpene composition, with monoterpene-linked sensory attributes providing the strongest and most consistent chemical-sensory associations within the RxCS population.

## Discussion

This study demonstrates that genetic variation at the CHR10_5431663 locus exerts a strong influence on monoterpene composition and relative abundance in wines produced from the Riesling × Cabernet Sauvignon population. This chemical variation translated, albeit imperfectly, into detectable differences in wine aroma. While monoterpenes were the primary drivers of genotype-dependent variation in volatile composition, their effects on sensory perception were modulated by additional matrix factors, resulting in continuous and partially overlapping phenotypes rather than discrete sensory classes.

Differences in wine volatile composition were most pronounced for monoterpenes, as indicated by both multivariate analyses and PERMANOVA results. Separation of wines along the primary axis of monoterpene PCA space was largely driven by (3*S*)-linalool and related compounds, underscoring the dominant role of (3*S*)-linalool in shaping wine monoterpene composition^9^ and consistent with the known role of monoterpene synthase genes located within the Chr. 10 genomic region.^13,33^ The differentiation observed between marker groups was expressed as a continuous gradient in monoterpene abundance rather than discrete classes, a pattern observed in previous studies.^12,21,34,35^ This suggests that variation in monoterpene composition reflects quantitative differences in biosynthesis, glycoside hydrolysis, and interconversion during fermentation, likely influenced by precursor availability and molecular rearrangement.^36–40^

In addition to total monoterpene abundance, substantial variation was observed in monoterpene composition among genotypes in this study. A meta-analysis of 19 studies identified 22 distinct monoterpenes occurring in grape and wine with (3*S*)-linalool and its derivatives constituting half of the compounds.^9^ Consistent with this finding, (3*S*)-linalool was the predominant compound across all RxCS wines regardless of monoterpene abundance, contributing between 22.6% and 79.7% of the total monoterpene pool. The identity and relative abundance of secondary monoterpenes varied considerably with compounds such as β-citronellol, *p*-cymene, and hotrienol frequently occurring as the second most abundant. Moreover, (3*S*)-linalool contributed a greater proportion of total monoterpenes in high-abundance genotypes. This pattern is consistent with the Chr. 10 locus acting primarily on (3*S*)-linalool biosynthesis rather than on downstream interconversion pathways.

In contrast to monoterpenes, other volatile compound classes exhibited weaker and less structured patterns of variation across wines. Esters, alcohols, and acids showed more diffuse distributions. Consistent with previous studies, phenylpropanoid pathway products such as 2-phenylethanol and phenylethyl acetate were considerably more abundant in Cabernet Sauvignon and a few progeny genotypes.^17,22,41^ Similarly, norisoprenoid content varied across genotypes, with Riesling and genotypes 28 and 54 exhibiting elevated levels of the characteristic Riesling aroma compound 1,1,6-trimethyl-1,2-dihydronaphthalene (TDN), consistent with genetic regulation of norisoprenoid accumulation in this population.

Comparison of berry and wine metabolite profiles revealed direct relationships primarily among monoterpenes. Although berry monoterpenes broadly correlated with those in wine, only (3S)-linalool and nerol showed significant berry-to-wine correlations, indicating that their abundance in wine is directly related to berry composition. Most berry monoterpene glycosides were highly correlated with a range of wine monoterpenes, yet *in vitro* enzymatic hydrolysis of Riesling monoterpene glycosides yielded a single monoterpene aglycone per glycoside,^42^ suggesting that glycoside hydrolysis alone does not account for the diversity of wine monoterpenes observed. Isomerization, cyclization, and other mechanisms of molecular rearrangement could explain the lack of direct correspondence between berry compounds and wine monoterpenes.^43^ Notably, two monoterpene glycosides (PH1 and PH2) correlated specifically with α-terpinene in wine, a compound that could not be linked to any berry monoterpene in our previous study.^13^ Surprisingly, no significant correlations between berry C6 alcohols and hexyl acetate were detected in the wines of this study despite previous documentation of acetate esters increasing in proportion with their corresponding precursor.^44^ While berry composition provides an important baseline, it is not solely determinative of wine volatile profiles.

The sensory consequences of genotype-dependent variation in monoterpenes were evident but not absolute. Descriptive analysis revealed significant product effects, with wines varying along gradients defined by attributes such as Floral, Tropical Fruit, and Apricot/Peach. These were strongly associated with monoterpene abundance in MFA. The association between (3*S*)-linalool and floral/fruit descriptors is well-established.^2,45–47^ Wines produced from CT genotypes tended to score higher for these descriptors, consistent with their generally elevated monoterpene levels. However, substantial overlap among genotypes was observed, and not all high-monoterpene wines exhibited correspondingly high sensory scores. For example, genotype 145, despite possessing relatively high monoterpene levels, was not strongly associated with monoterpene-derived sensory attributes. This decoupling highlights the complexity of aroma perception, which depends not only on compound concentration but also on odor thresholds, interactions among volatiles, and matrix effects.

Despite identical fermentation conditions and obscuration of wine color during evaluation, the phenolic matrix associated with berry pigmentation accounted for much of the variation among wines in this study. Regardless of monoterpene level, pigmented genotypes grouped more closely with each other than with unpigmented individuals of comparable monoterpene abundance, indicating that matrix composition strongly influences aroma expression. Differences in genotype-dependent separation between red and white wines further support this interpretation: clearer gradients were observed among red wines, whereas white wines exhibited weaker structure and greater overlap among marker classes. Although wine color was not visible to panelists, odor alone has been shown to be sufficient to distinguish red from white wines,^48^ suggesting that color-associated aroma differences may have contributed to separation in the sensory data independent of monoterpene content. However, the white wines in this study were atypical in that they were fermented on the skins, and matrix effects from elevated phenolic content likely modulated aroma expression in both color groups. Further work involving a larger number of genotypes within a single color group could help clarify the contribution of (3*S*)-linalool to wine aroma perception.

Matrix effects are known to modulate the perception of volatile compounds in wine.^49–53^ Ethanol suppresses the volatilization of polar aroma compounds while enhancing that of nonpolar ones, and has been shown to alter the perception of aroma descriptors.^54–56^ Phenolic compounds such as gallic acid, catechin, and quercetin can inhibit both the release of monoterpenes from glycosidic precursors and their subsequent volatilization through intermolecular interactions, including hydrogen bonding and dispersive forces.^57,58^ (3*S*)-Linalool, in particular, has been shown to interact strongly with phenolic acids, which may reduce its volatility and perceived intensity. In the context of this study, these interactions may have attenuated the perceived intensity of (3*S*)-linalool-associated aromas in wines produced from pigmented genotypes without completely eliminating relative differences among genotypes, resulting in lower sensory scores relative to white wines while preserving genotype-dependent separation. Additionally, our decision to ferment all genotypes on their skins may have contributed to the lack of differentiation between white genotypes by increasing phenolic content, thereby partially obscuring genotype-dependent differences in monoterpene-associated aroma.

In addition to chemical interactions, aroma perception is influenced by the mode of sensory evaluation. Wine aroma is perceived through both orthonasal and retronasal pathways, which can differ in their sensitivity to matrix effects.^59^ While this study focused on orthonasal evaluation, retronasal perception plays a critical role in wine aroma perception, and interactions between aroma compounds, the wine matrix, and oral components such as mucosal surfaces can further alter volatility, persistence, and perception,^60,61^ Together, these factors suggest that matrix-mediated modulation of aroma release and perception may partially mask genotype-dependent chemical differences.

Integration of chemical and sensory data through MFA further highlighted the complexity of relationships between volatile compounds and aroma attributes. While strong associations were observed between monoterpenes and floral and fruity descriptors, many expected relationships, particularly between esters and red- or dark-fruit-related attributes, were weak or absent.^8,62,63^ This finding is consistent with previous work showing that the sensory impact of individual compounds in wine cannot be reliably inferred from their concentrations alone.^31^ Instead, aroma perception emerges from complex interactions among multiple volatiles, including additive, synergistic, and suppressive effects.^4,55,64^ For example, the presence of (3*S*)-linalool has been found to reduce the sensory impact of acetate esters and alter the perception of limonene.^10,47^ The lack of strong associations between yeast-derived esters and fruit descriptors suggests that these compounds may contribute to background aroma or interact with other volatiles in ways not captured by simple correlation analyses. The absence of “petrol” or “diesel” from the consensus descriptor list is notable, given that TDN is present in all wines. Although TDN concentration increases during wine aging, peri- or suprathreshold levels are common in Riesling wines less than two years old,^65^ and concentrations in the RxCS wines may still have been insufficient to surpass the individual identification threshold.

In this study, we evaluated only wine aroma characteristics; however, the results have broader implications for wine grape breeding. The strong and consistent effect of the CHR10 locus on monoterpene composition indicates that CHR10_5431663 could serve as an effective marker for selecting enhanced floral and fruity aroma potential in breeding programs. At the same time, the incomplete translation of chemical differences into sensory outcomes highlights the need to consider the broader metabolic and matrix context when selecting for aroma traits. Selection based solely on a single compound or locus may not yield predictable sensory profiles, particularly in complex matrices such as wine. Even within the limited set of 24 genotypes examined, substantial variation was observed in ripening time, pH, titratable acidity, yield, and Brix. These traits are of clear importance for the development of new cultivars adapted to current and future viticultural challenges.^66^ For example, warmer temperatures, particularly elevated nighttime temperatures, can accelerate the loss of acidity and anthocyanins in grapes.^67–69^ Understanding the sensory consequences of variation in aroma-related pathways is therefore critical, as alleles associated with desirable agronomic traits may also carry linked effects on aroma composition, ultimately influencing the sensory expression of newly developed cultivars.

In conclusion, this study demonstrates that genotype-dependent variation in wine aroma arises from coordinated changes in monoterpene composition driven by allelic variation at a key genomic locus, with additional modulation by other volatile compounds and matrix effects. The integration of genetic, chemical, and sensory data reveals a clear but non-linear relationship between genotype and phenotype, underscoring the complexity of aroma formation in wine and the importance of a systems-level approach to its study.

## Acknowledgements

The project was partially funded by the Ray Rossi Endowment in Viticulture and Enology and the E.J. Gallo Winery. JL was partially supported by the American Society for Enology and Viticulture; the American Wine Society Educational Foundation; the Horace O. Lanza, Margrit Mondavi, Harold P. Olmo, Pearl & Albert J. Winkler, Robert Lawrence Blazer, Louis R. Gomberg, Knights of the Vine, and the Wine Spectator scholarships; and the Albert G. and Lilian R. Fradkin Student Award Fund.

